# A critical reexamination of recovered SARS-CoV-2 sequencing data

**DOI:** 10.1101/2024.02.15.580500

**Authors:** F. Débarre, Z. Hensel

## Abstract

SARS-CoV-2 genomes collected at the onset of the Covid-19 pandemic are valuable because they could help understand how the virus entered the human population. In 2021, Jesse Bloom reported on the recovery of a dataset of raw sequencing reads that had been removed from the NCBI SRA database at the request of the data generators, a scientific team at Wuhan University (Wang *et al*., 2020b). Bloom concluded that the data deletion had obfuscated the origin of SARS-CoV-2 and suggested that deletion may have been requested to comply with a government order; further, he questioned reported sample collection dates on and after January 30, 2020. Here, we show that sample collection dates were published in 2020 by Wang *et al*. together with the sequencing reads, and match the dates given by the authors in 2021. Collection dates of January 30, 2020 were manually removed by Bloom during his analysis of the data. We examine mutations in these sequences and confirm that they are entirely consistent with the previously known genetic diversity of SARS-CoV-2 of late January 2020. Finally, we explain how an apparent phylogenetic rooting paradox described by Bloom was resolved by subsequent analysis. Our reanalysis demonstrates that there was no basis to question the sample collection dates published by Wang *et al*..

**Note for bioRxiv readers:** The automatically generated Full Text version of our manuscript is missing footnotes; they are available in the PDF version.

## 1. Introduction

In June 2021, Jesse Bloom described the recovery of SARS-CoV-2 sequencing read data that had been deleted from the Sequence Read Archive (SRA) at the request of the data generators based at Wuhan University (Bloom, 2021a). Bloom claimed that the recovered data shed light on the early days—and thereby the origin—of the Covid-19 pandemic. His results, initially presented in a bioRxiv preprint and accompanied by a Twitter thread, reverberated in popular media^1,2^ and were addressed at a press conference by a vice minister of China’s National Health Commission.^3^ Bloom’s study was later published in *Molecular Biology and Evolution* (MBE; Bloom, 2021b).

The study for which the sequencing data had been generated, suddenly under international public scrutiny, presented a diagnostic technique based on amplifying fragments from a portion of the SARS-CoV-2 genome using nanopore technology (Wang *et al*., 2020b). The article, published in the journal *Small*, had initially been shared as a preprint on the medRxiv server (Wang *et al*., 2020a) (see Table 1 for a timeline). After the preprint was posted, raw sequence read data were submitted by Wang *et al*. to SRA as Bio-Project PRJNA612766 in mid-March 2020; these data were removed in mid-June 2020. Neither the preprint nor the published article mentioned the public availability of raw sequencing data.

**Table 1:**
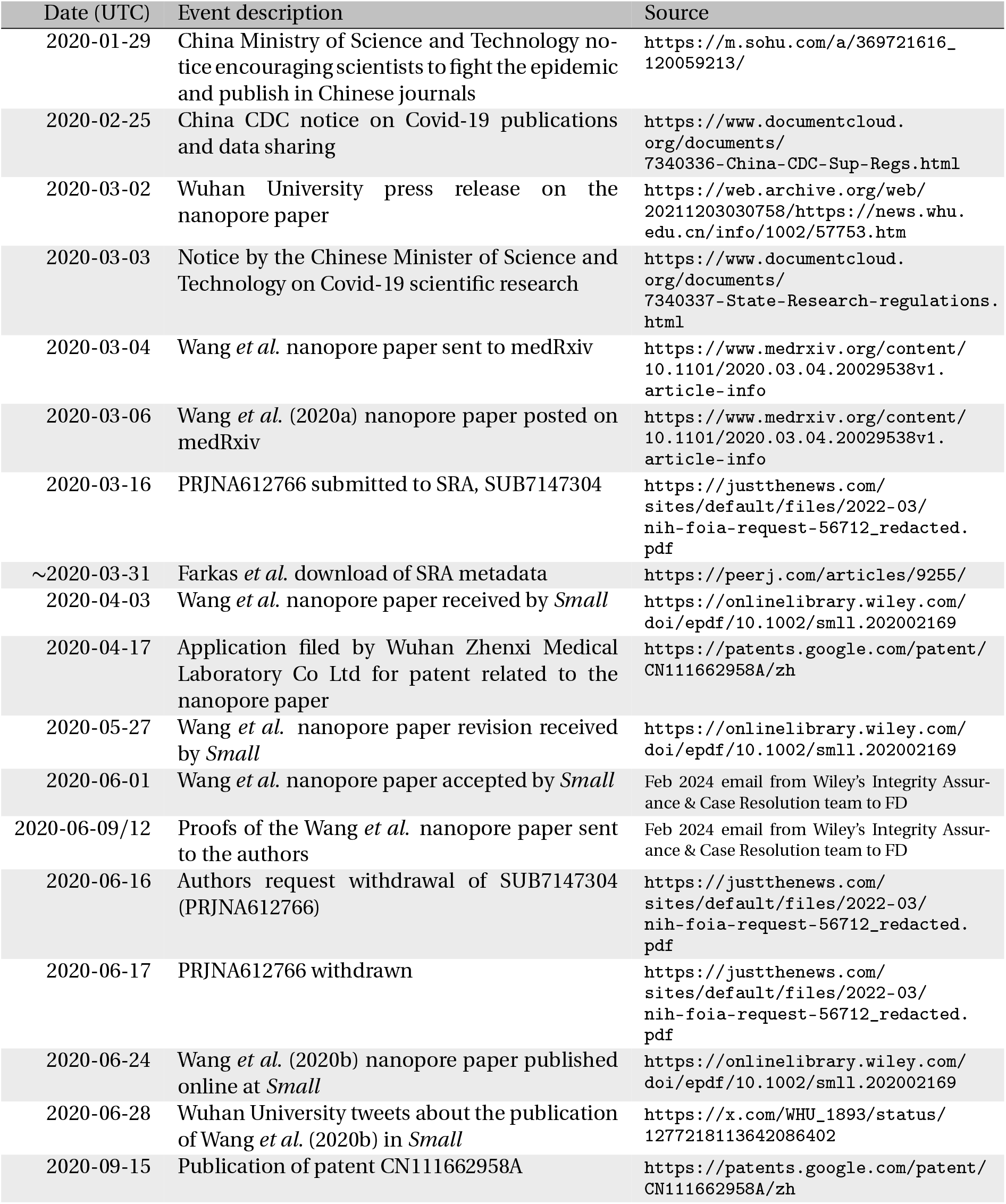
Timeline of 2020 events related to Wang *et al*.‘s study and sequencing data. The dates are written in the YYYY-MM-DD format.

Central to Bloom’s claim was the argument that the removal of these data by Chinese scientists had “*obscured sequences relevant to the early spread of SARS-CoV-2 in Wuhan*” (Bloom, 2021b). The argument is part of a narrative, presented in Bloom’s introduction, which claims that Chinese researchers were silenced by China’s government, and had to retract previously released data related to cases prior to mid-December 2019 to comply with one government order or another.^4^ Wang et al. however submitted their raw data to SRA weeks *after* the government orders (see Table 1), contradicting the censorship narrative.

Although the raw sequencing data had been removed from the SRA by the National Center for Biotechnology Information (NCBI) after a request by the authors, the published, peerreviewed article (Wang *et al*., 2020b) included as its Table 1 all of the mutations identified in the raw reads, the same ones later found in Bloom’s reanalysis. The preprint (Wang *et al*., 2020a) contained a less complete version of the table that nevertheless identified the mutation central in Bloom’s analysis: mutation C29095T, in sample C2. Bloom (2021b)’s reanalysis indicated that Wang *et al*.‘s results were consistent with data they submitted to SRA. In other words, the information contained in the raw data that Bloom recovered from SRA was available and described in manuscripts published in 2020.

Bloom initially discovered the existence of the Wang *et al*. sequences via a paper which had referenced sequencing data published on the SRA at the end of March 2020 (Farkas *et al*., 2020). Sequencing data relating to Wang *et al*. were listed in Supplementary Table 1 of Farkas *et al*. (2020), but the data were no longer available and not findable on SRA when Bloom looked for them in 2021. However, the data had been backed up to the cloud, and Bloom (2021a) recovered most sequencing data from the backup. Bloom (2021b) obtained the remaining sequencing data from a March 2020 backup made available by Lifebit Life Sciences that included raw sequence data and metadata for all 241 runs listed in Farkas *et al*. (2020).^5^

In reply to Bloom’s preprint, in 2021, Wang *et al*. responded that the samples from which sequences had been obtained had been collected on 30 January 2020 at the earliest.^6,7^According to the authors, the partial sequences were therefore not relevant to the origin of SARS-CoV-2. A co-author of Wang *et al*. also described the rationale for withdrawal of sequencing data from the SRA in an interview:^8^ raw sequence data had been submitted to the SRA to accompany the submitted manuscript. After their article was accepted by the journal *Small*, the proofs received by the authors did not include a data availability statement. The authors thought it was appropriate to request deletion because journal editors did not retain their data availability statement and because SRA data would not be referenced in the manuscript.^9,10^ This explanation is consistent with the timeline of events (see Table 1): the request for data removal took place after the authors had been sent the proofs of their article, and before their article was online. Wiley’s Integrity Assurance & Case Resolution team, who conducted at our request an investigation on the case, confirmed that the data availability statement was removed by the journal during copy-editing.^11^

In his study, Jesse Bloom explored possible roots of the early SARS-CoV-2 phylogeny in the context of the data from Wang *et al*.. Assuming that the sequence of the most recent common ancestor of SARS-CoV-2 should resemble the most closely related viruses sampled in bats, and neglecting sampling dates, Bloom considered three roots: all were of lineage A (defined by substitutions C8782T and T28144C compared to the reference sequence Wuhan-Hu-1), with each proposed root haplotype containing one additional mutation towards a bat-virus outgroup (either T3171C, C18060T, or C29095T) compared to Wuhan-Hu-1 (which is of lineage B; the positions and names are summarized in Table 2). This analysis also did not account for the fact that C → T is the most frequent type of single-nucleotide substitution in SARS-CoV-2 genomes (Azgari *et al*., 2021; De Maio *et al*., 2021; Bloom *et al*., 2023; Ruis *et al*., 2023).

**Table 2:** Substitutions in the different lineages and Bloom’s proposed roots. We use Wuhan-Hu-1 as reference. The positions highlighted with a star (^*\**^) are covered in the “recovered” sequences. The lineage defined by A + C18060T is referred to as “proCoV2” by Bloom (2021b), following previous analysis (Kumar *et al*., 2021). Since the original “proCoV2” had three additional substitutions in the Kumar *et al*. preprint, we avoid this nomenclature.

| Position | 3171 | 8782 | 18060 | 28144* | 29095* |
| --- | --- | --- | --- | --- | --- |
| Lineage B (Wuhan-Hu-1) | T | C | C | T | C |
| Lineage A | T | **T** | C | **C** | C |
| Bloom 1: A + C18060T | T | **T** | **T** | **C** | C |
| Bloom 2: A + C29095T | T | **T** | C | **C** | **T** |
| Bloom 3: A + T3171C | **C** | **T** | C | **C** | C |

Rooting the SARS-CoV-2 tree had long been identified as a difficult problem (Pipes *et al*., 2021), for which different methods give different answers (Pekar *et al*., 2021). In particular, an early sequence with three spurious mutations caused rooting issues (J. Wertheim, personal communication; Pekar *et al*., 2022) until these errors were corrected in the China-WHO joint mission report (World Health Organization, 2021, Table 6, ID: S02, IPBCAMS-WH-01). Pekar *et al*. (2022) later showed that the root almost certainly lies along one branch including lineages B and A. However, uncertainty remains regarding whether the ancestral state is lineage A, lineage B, or an intermediate between them.

Bloom’s rooting methodology led to a known conundrum (Rambaut *et al*., 2020): all three of the roots that Bloom considered plausible roots did not resemble the sequences with earliest collection dates, defying the expectation “*that [the outbreak’s progenitor] should be among the earliest sequences*”. To explain this discrepancy, Bloom suggested that critical early (meta)data may be missing or altered, identifying the “recovered sequences” as examples for which true collection dates were potentially earlier than reported: “*The press conference and blog posts also stated that the sequences were all collected on or after January 30, 2020, rather than “early in the epidemic” as originally described in Wang et al. (2020)*.”.

Here we provide multiple lines of evidence showing that the 30 January 2020 collection date was correct – including the crucial facts that collection dates were available in datasets analyzed by Bloom, and that the 30 January 2020 date was manually removed by Bloom during his analysis. Speculation that the samples reported by Wang *et al*. were some of the earliest samples from Covid-19 patients implies that reported collection dates are false. This speculation was, and remains, unsubstantiated and unwarranted.

## 2. Results

### 2.1 The 30 January 2020 collection date was present in the data from Wang *et al*. (2020), but manually removed by Bloom during his analysis of the data

In his article, Bloom questioned the veracity of 30 January 2020 collection dates reported by authors of Wang *et al*. (2020b) in 2021. However, there is contemporaneous evidence confirming the dates. The collection dates were indeed present in the SRA metadata and remain visible today.^12^ The collection date visible today, 30 January 2020, matches the date reported by authors of Wang *et al*. in 2021. According to the SRA team, the collection dates were the same in 2020. This is further confirmed by Supplementary Table 1 of Farkas *et al*. (2020), which was compiled at the end of March 2020 (see Figure 1A). The same supplementary table was used by Bloom (2021a,b) to originally identify raw sequencing data from Wang *et al*. (2020b). This table consists of sequencing metadata downloaded after publication of the Wang *et al*. (2020a) preprint, but before its submission to *Small*; see Table 1 for a chronology. In summary, there is zero evidence that co-authors of Wang *et al*. ever fabricated or altered sample collection dates, and ample evidence that they did not. The 30 January 2020 collection date was known to Bloom and then was manually deleted and replaced by “early in epidemic” (see Figure 1B). Bloom did not modify February collection dates. In addition to the supplementary table from Farkas *et al*. (2020), matching collection dates were again available in metadata from Lifebit Life Sciences, from a direct back-up of (meta)data from the SRA.

**Figure 1:**
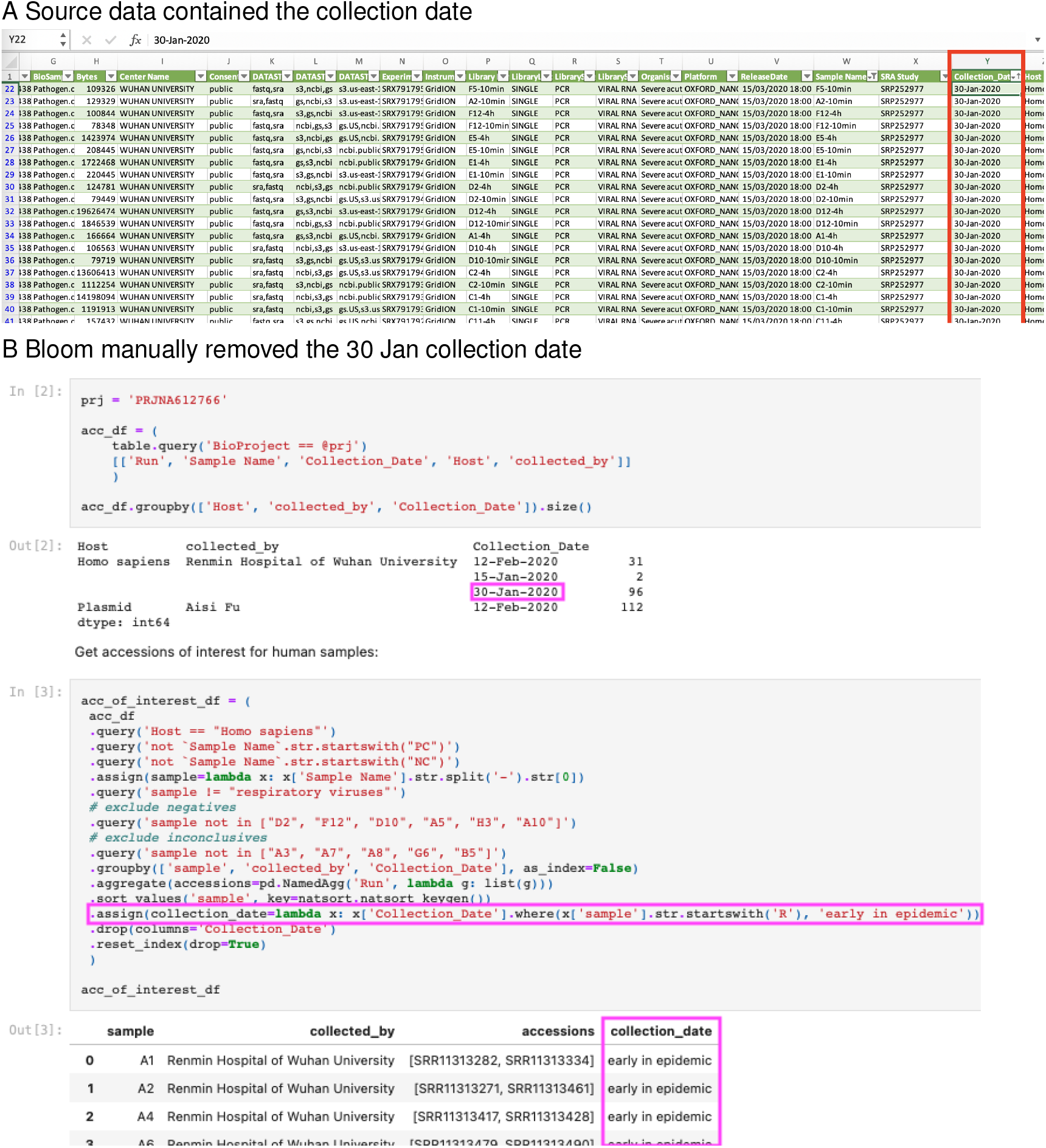
Presence and removal of the 30 Jan collection date. A: Screenshot of Supplementary Table 1 from Farkas *et al*. (2020). The highlighted cell is a collection date (30-Jan-2020). Source: https://dfzljdn9uc3pi.cloudfront.net/2020/9255/1/Supplementary_Table_1.xlsx; https://peerj.com/articles/9255/#supp-2. The file is also available in Jesse Bloom’s Github repository (https://github.com/jbloom/SARS-CoV-2_PRJNA612766/blob/main/manual_analyses/PRJNA612766/Supplementary_Table_1.xlsx). B: Screenshot of Bloom’s code in the `manual_analyses` folder. Highlighted are the collection date, initially correctly loaded (top); the code line removing it (middle); and the result (bottom). Source: https://github.com/jbloom/SARS-CoV-2_PRJNA612766/blob/main/manual_analyses/PRJNA612766/extract_accessions.ipynb. The date was modified in git commit `92aaae9`.

Collection dates on and after 30 January 2020 are significant, in that a small number of partial sequences from samples collected at this time are unlikely to substantially shift likelihoods of proposed SARS-CoV-2 progenitor genomes. Full genome sequences from samples collected on or before 30 January are not rare: there were 507 such sequences in data considered by Bloom (2021b) (including 365 from China). Our analyses use Bloom (2021b)’s dataset, and we additionally check that conclusions still hold with a second dataset obtained after more stringent quality control (Pekar *et al*., 2022) and the addition of recently published sequences (Lv *et al*., 2024), containing 448 sequences collected on or before 30 January 2020, 318 of which from China.

### 2.2 The study timeline is incompatible with the alleged censorship

Bloom speculated about a scenario in which co-authors of Wang *et al*. were compelled to remove their data from SRA under the pressure from China’s government, which published notices regarding scientific publication during the Covid-19 pandemic. The actual timeline of events contradicts this narrative: the preprint itself was posted to medRxiv *after* the publication of the second notice referenced by Bloom, as were the raw sequencing data on the SRA (see Table 1). The notices did not prevent Wang *et al*. from sharing information.

As acknowledged by Bloom (2021b), information on the mutations contained in Wang *et al*.‘s sequences was not removed, but contained in a Table in their article. The fact that Wang *et al*.‘s description of the mutations identified in their samples is *even more complete* in the published article than in their preprint (Wang *et al*., 2020a,b, Table 1) directly refutes the hypothesis that the raw sequence reads were removed from the SRA to obfuscate the origin of SARS-CoV-2.

### 2.3 The “recovered” sequences are compatible with a late January collection date

To further test the veracity of the 30 January 2020 sampling date announced by Wang *et al*., we compare partial sequences from Wang *et al*. (2020b) to the corresponding region of other early sequences, following the approach in Figure 4 of Bloom (2021b).

As a first comparison, we turn to sequencing data generated via a similar nanopore-based technology, obtained from samples collected in Wuhan, with similar collection dates to those reported for the earliest samples in Wang *et al*.. These data were reported in the context of an article by Yan *et al*. (2021); the samples were collected from “*various Wuhan health care facilities*” on 25 and 26 January 2020, and consensus sequences were deposited on GISAID. These data were contained in the dataset analyzed by Bloom (2021b). Two sequences from the Yan *et al*. (2021) dataset are present in proposed progenitor nodes in Bloom (2021b).^13^

Figure 2 shows that the distribution of substitutions in sequences from Yan *et al*. (2021) is similar to that of Wang *et al*. (2020b) (“recovered” sequences). In particular, the proportions of the lineage-A defining mutation T28144C are indistinguishable in the two datasets (17/42 in the Yan *et al*. (2021) data contained in Bloom‘s dataset and 5/13 in the “recovered” sequences; Fisher’s Exact Test, *p* = 1); so are also the proportions of the C29095T mutation highlighted by Bloom (1/42 in the Yan *et al*. (2021) data and 1/13 in the “recovered” sequences; Fisher’s Exact Test, *p* = 0.42). The two distributions remain similar when the outgroup comparator is changed (Figure S1) or when a partial dataset of Yan *et al*. (2021) is used, in which sequences with potential sequencing errors were removed (Figure S2).

**Figure 2:**
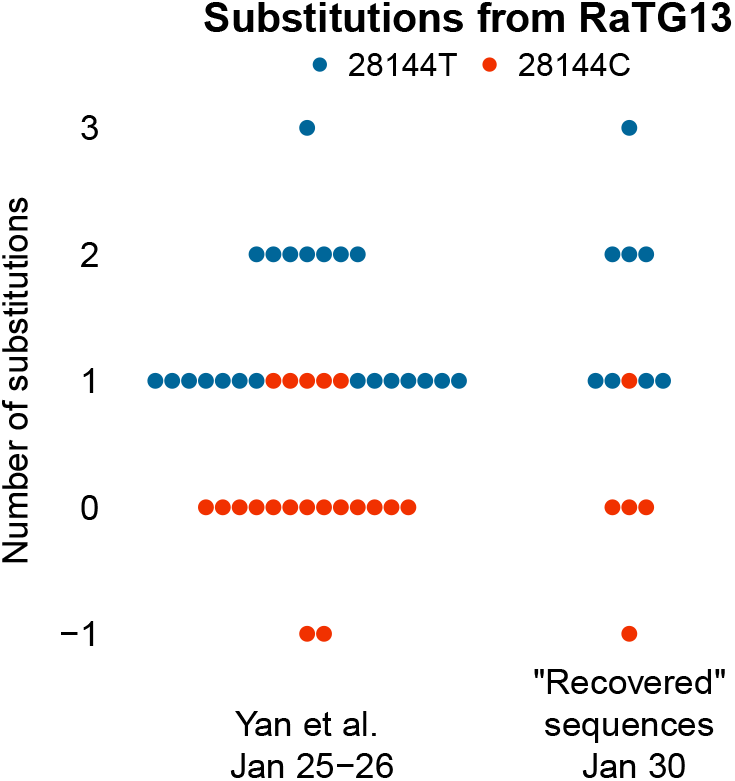
Number of substitutions from bat SARS-like coronavirus RaTG13 in the region between nucleotides 21,570–29,550 that is considered in Figure 4 in Bloom (2021b) (relative to lineage A, which is equivalent to lineage A + C18060T [“proCoV2”] in this region, as position 18060 is not covered). Sequences from Yan *et al*. (2021) are compared to those from Wang *et al*. (2020b) (“Recovered” sequences). Substitutions are counted such that 0 corresponds to the same distance as between RaTG13 and lineage A; negative values (*−*1) correspond to additional substitutions towards RaTG13 (C29095T for a “recovered” sequence and for one of the Yan *et al*. (2021) sequences, and C22747T for the other Yan *et al*. sequence). Substitution T28144C is characteristic of lineage A and is highlighted in red. (NB: We use RaTG13 only for the sake of comparison with Bloom’s analysis.)

Figure 2 also illustrates that substitutions towards the chosen outgroup are not necessarily signs of their ancestral nature. The −1 positions of three sequences in Figure 2 are due to C29095T (one “recovered sequence” and one sequence from Yan *et al*. (2021)) and to C22747T (the other Yan *et al*. (2021) sequence). Both substitutions have subsequently reappeared in other SARS-CoV-2 lineages (see Figure S3). Outside of the region covered in sequences from Wang *et al*., the Yan *et al*. sequence with C22747T also contains T4402C and G5062T, identifying C22747T as a reversion subsequent to mutations that characterize a common early epidemic genome in lineage A sampled in China (Beijing), South Korea, and Japan (i.e., not an early ancestral genome).

The comparison can be extended to a broader set of early sequences. To do so, we used Pekar *et al*. (2022)’s carefully curated set of early sequences, and complemented it with recently published early sequences from Shanghai (Lv *et al*., 2024) to build a comprehensive set of sequences available to date. Figure 3 shows that the number of substitutions in the Wang *et al*. (2020b) dataset is consistent with those observed in other sequences with similar collection dates. The pattern holds when changing the comparator (Figure S4; all data points instead of averages are shown in Figure S5).

**Figure 3:**
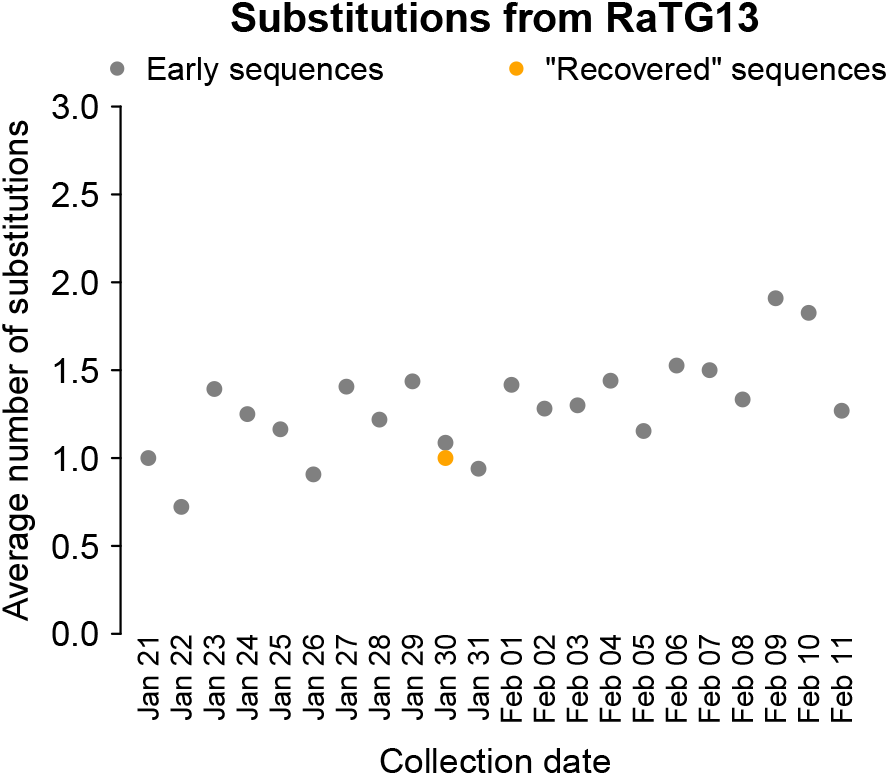
Average number of substitutions from bat SARS-like coronavirus RaTG13 (relative to lineage A, or, equivalently, lineage A + C18060T), over SARS-CoV-2 nucleotides 21,570–29,550, comparing all available sequences (after de-duplications and curation (Pekar *et al*., 2022), and addition of recently published sequences (Lv *et al*., 2024); gray points) to Wang *et al*. (2020b)’s “recovered” sequences (orange).

The same conclusion could readily be obtained with the set of sequences studied by Bloom (2021b) (see Figure S6). Focusing on sequences collected ±7 days around 30 January 2020, the proportions of key mutations such as T28144C and C29095T are, again, indistinguishable between Bloom‘s dataset and the recovered sequences (T28144C: 226/650 in Bloom (2021b)’s dataset; Fisher’s Exact Test, *p* = 0.78; C29095T: 30/650; Fisher’s Exact Test, *p* = 0.47). Given the proportion of C29095T in available sequences (4.6% in Bloom’s dataset), there was a high likelihood of finding this specific mutation in at least one of 13 additional sequences. C29095T is only one of many early-pandemic mutations towards RaTG13 that could have been observed in these samples. The “recovered” sequences dataset is therefore unremarkable; it is consistent with expectations for samples collected in Wuhan on 30 January 2020.

Importantly, by implicitly assuming that positions that are not covered are not mutated, Bloom‘s methodology will underestimate divergence for sequences with low coverage. Bloom (2021b) highlighted “*a sequence (Guangdong/FS-[S]30-P00502/2020 reportedly collected in late February that is actually two mutations more similar to RaTG13 than lineage A + C18060T* “ (corresponding to a point at “ −2” in the “Other China” panel of his Fig. 2). We doubt it is a coincidence that the most striking outlier in this figure is also a sequence with one of the lowest levels of coverage in his dataset (84%, ranking 5th of 1886 sequences). Absence of coverage is not equivalent to absence of mutation.

The “recovered” sequences’ partial coverage also implies that there is no evidence that any of them actually matched Bloom’s proposed roots. First, key positions are missing. In particular, it was misleading to define mutations relative to A+C18060T (called “proCoV2”) instead of simply relative to A in Bloom (2021b)’s Table 1: writing that there are no substitutions relative to A+C18060T invites the reader to believe that some of the “recovered” sequences matched the A+C18060T root, while the 18060 position is simply not covered. Secondly, because the sequences are partial, there is no guarantee that there are not additional mutations in the large parts of the genome that were not covered, that would move the sequences away from the proposed roots.

### 2.4 No evidence of a widespread undetected circulation of virus with C29095T in Wuhan

Analysis in Bloom (2021b) highlighted sequences collected in the Guangdong province that were related to what he considers a very plausible progenitor SARS-CoV-2 genome (lineage A + C29095T). The presence of C29095T brought them closer to the bat virus outgroup, and positioned them at a striking “−1” in Figure 4 of Bloom (2021b) (orange dots in his figure). Initially described as belonging to “*two different clusters of patients who traveled to Wuhan in late December of 2019*”, these sequences could be interpreted as evidence of a widespread but so far undetected circulation of similar viruses in Wuhan in late 2019.

Examination of the included sequences, however, indicated that there was only one cluster rather than two; Bloom recently corrected the article after we and others pointed this out (Bloom, 2023). All the patients were from the same family group, and therefore the sequences were not independent. In addition, we found that multiple sequences collected from the same patients were included in Bloom’s dataset, sometimes labeled as “Other China”. Briefly, at least seven sequences belong to the same family cluster detected in Guangdong (Chan *et al*., 2020; Kang *et al*., 2020), corresponding to four patients, two of which were sampled multiple times; two of the sequences were labeled as “Other China” by Bloom (see supplementary Tables S1 and S2 for details). Further, following Bloom (2021b)’s advice to “*[go] beyond the annotations in GISAID to carefully trace the location of patient infection and sample sequencing*”, we note that plausible index patients in the Guangdong cluster did not just travel to Wuhan in late December 2019, but had visited a relative hospitalized in Wuhan for febrile pneumonia (Chan *et al*., 2020). In other words, they had been to one of the few places other than the Huanan market where one was most likely to encounter other people infected by SARS-CoV-2 in Wuhan at that early date.

Bloom argued that a A + C29095T root is “*now more consistent with the evidence that the pandemic originated in Wuhan, as half*^14^ *its progenitor node is derived from early Wuhan infections, which is more than any other equivalently large node*.”. There was in fact a single “recovered” sequence in the node (sample C2); the rest of the increase in support was due to the relabeling of Guangdong cluster sequences. Moreover, in addition to the documented epidemiological link between annotated cases discussed above, there is no reason to expect that sample C2 necessarily lacks additional mutations outside of the region covered in “recovered” sequences.^15^

Furthermore, epidemiological links to Wuhan are very common in case reports from January 2020, and not only for A + C29095T. For example, all eight sequences in Bloom’s proposed A + T3171C root have a documented epidemiological link to Wuhan (Jiang *et al*., 2020), as does the first Covid-19 case detected in the United States with A + C18060 (Holshue *et al*., 2020). But this remark should not be interpreted as support for those haplotypes as roots, because lineage A and lineage B exports were for instance found in Australia (Eden *et al*., 2020) and in the first two cases in Thailand (Okada *et al*., 2020). Although it is difficult to systematically explore the epidemiological history of all early pandemic sequences, a recent report with epidemiological history for a large number of patients in Shanghai makes this possible (Lv *et al*., 2024). Epidemiological history is described for 13 of 14 patients with samples collected on or before 31st January 2020 (the same period used for phylogenetic analysis in Bloom’s study); 8 have direct links to Wuhan via travel history. Lastly, a complete annotation of exposure history for cases outside of Wuhan should note the case with symptom onset predating any case in the Guangdong cluster by almost two weeks: this is a lineage B case identified in Beijing with a link not only to Wuhan, but to the Huanan market specifically (Liu, 2020).

The history of the Guangdong cluster indicated that the C29095T substitution was present in Wuhan in late December 2019; it is therefore unsurprising that it was sampled in late January 2020 as well. While C29095T is a mutation towards the most closely related viruses sampled in bats, the A + C29095T haplotype is rejected as the root of the SARS-CoV-2 phylogenetic tree in humans by Pekar *et al*. (2022), and confirmed by more recent analyses including recently published sequences (FD & ZH, personal communication). Methods in Bloom (2021b) neglect that C → T mutations are by far the most frequent type of mutation during the pandemic (Azgari *et al*., 2021; De Maio *et al*., 2021; Bloom *et al*., 2023; Ruis *et al*., 2023), and that the C29095T mutation, specifically, occurs much more frequently than expected for a typical C*→*T mutation (Bloom and Neher, 2023, supplementary data `nt_fitness.csv`). This mutation regularly reappeared in multiple lineages during the last four years of SARS-CoV-2 evolution in humans (Figure S3). For example, it is a defining mutation in the HP.1.1 lineage that emerged in North America in mid-2023. In fact, C29095T is even recurrent in Bloom (2021b)’s phylogenetic trees (Figs 3 and 5), where this position mutates three times.

## 3. Discussion

The facts we present do not support the conclusion that recovered sequences “shed more light on the early Wuhan SARS-CoV-2 epidemic” as promised by Bloom (2021b). First, Bloom’s phylogenetic analyses did not require any of the recovered raw data, as all the data utilized were publicly available in a peer-reviewed article (Wang *et al*., 2020b). Second, the “recovered” sequences are partial: only a fraction of the whole genome was sequenced, by design, seriously limiting the usefulness of these “recovered” sequences to infer ancestral states. Finally, the samples were collected in late January 2020, and as such were unlikely to provide useful information on the genome of the proximal ancestor of SARS-CoV-2 prior to the outbreak in Wuhan. Mutations identified in these samples, including C29095T, are unsurprising to find in Wuhan in late January.

The haplotypes proposed by Bloom as roots of the SARS-CoV-2 tree led to a conundrum: they were of lineage A, while the earliest known sequences were of lineage B, as were all the Huanan market sequences known at the time. The issue of the absence of lineage-A sequences directly linked to Huanan market was resolved a few months after the publication of Bloom’s article. In early 2022, Liu *et al*. (2022) revealed that a lineage-A genome had been detected in an environmental sample collected in the Huanan market on the 1st of January 2020. Raw sequencing data from this sample, shared a year later (Liu *et al*., 2024), confirmed the lineage assignment (Crits-Christoph *et al*., 2023).

The question of the precise identity of SARS-CoV-2’s root remains unresolved. Pekar *et al*. (2022) analyzed a SARS-CoV-2 origin scenario resolving the rooting conundrum: the root may never have been in humans, but only in the animals from which SARS-CoV-2 spilled over more than once. Under this scenario, the two early lineages, A and B, would have been the products of two spillovers close in time and space, possibly from the same group of animals. The more “bat-like” lineage A likely spilled over after B, resulting in most early sequences being derived from lineage B. Among the several examples of animal-to-human SARS-CoV-2 transmission documented later in the pandemic (reviewed in EFSA Panel on Animal Health and Welfare (AHAW) *et al*., 2023), such a scenario of multiple transmissions close in time and space, from a group of animals to humans, occurred notably with pet hamsters in Hong Kong, for which a genomic investigation indicated that different patients had been independently infected by hamsters at a pet shop (Yen *et al*., 2022). Low diversity in viral genomes identified in samples from bats at the same time and place is also common; for example, RshSTT182 and RshSTT200 genomes differ by only 3 nucleotides (Delaune *et al*., 2021).

Facing the same conundrum, Bloom proposed a different explanation: the record of published SARS-CoV-2 genomic sequences could have been altered by China authorities by selectively suppressing publication of sequences from the earliest samples from patients unlinked to Huanan market. We have demonstrated that Bloom’s analysis does not support this speculation.

A divergence between data and expectations from a theory can be due to issues with the data, or to issues with the theory. Checking the reliability of data, especially from diverse sources, is an essential step in any scientific endeavor. There are for instance some aberrant collection dates in public databases, and data need to be curated to avoid absurd conclusions due to issues in input data. When the data are consistent, however, it is also important to challenge one’s theory and to reconsider methodological choices. Hybrid methods taking into account both the bat virus relatives and collection dates find support for A or B roots, but confidently reject both the A + C29095T and A + C18060T roots considered by Bloom (2021b) (Pekar *et al*., 2022, Table 1).

In an attempt to reconcile the proposed roots with the lack of supporting data, Bloom (2021b) suggested that the collection date indicated by Wang *et al*. in 2021 could be incorrect. Our investigation finds no support for this suggestion. Contrary to Bloom‘s description, there was no contradiction in Wang *et al*. (2020b)’s statements on collection dates: the date they gave in 2021 in reply to Bloom’s preprint (30 January 2020) was the same as the date they had submitted to the SRA in early 2020. Further, the term “early in the epidemic” depends on its context. We have found examples of it being used to refer to very different periods. Notably, Bloom defines “Early SARS-CoV-2 Sequences” as those collected prior to March 2020. In addition, the meaning of “early in the epidemic” depends on the location: “early” in Brazil’s outbreak is later than in China, “early” in Shanghai is later than in Wuhan, and “early” at Renmin Hospital is later than at hospitals closer to the epicenter of the outbreak. The context of the study by Wang *et al*. is one in which fever clinics at Renmin Hospital were suddenly overwhelmed with demand for molecular testing of suspected Covid-19 one week after the beginning of Wuhan’s lockdown (Liu *et al*., 2020) and only a few days after Renmin hospital was reportedly still sending staff to supplement Jinyintan hospital where the earliest patients were sent for treatment.^16^

Thus, 30 January was indeed early in the context of the epidemic at Renmin hospital, and Wang *et al*.‘s use of the term in their preprint and peer-reviewed paper was therefore consistent with the 30 January collection date. Late January is, however, not so early that the “recovered” sequences can be considered as coming from the “earliest samples” ever collected from Covid-19 patients, yet this is what was widely reported as a conclusion of Bloom’s work.^17^ There was therefore no inconsistency when it was said that 30 January is not early in the context of “*Covid-19 origin tracing*”, at a press conference^18^ following the publication of Bloom’s preprint (Bloom, 2021a).

The unavailability of the Wang *et al*. raw sequencing data on the SRA after June 2020 was shown to be the product of a human error (Berman *et al*., 2022). According to policies of the International Nucleotide Sequence Database Collaboration (INSDC; Brunak *et al*., 2002), of which the SRA is a member, data once made public on this repository are supposed to belong to the scientific record, and to remain accessible. Mechanisms exist to remove data from indexing, but keep them available by accession (“suppress” command) (Berman *et al*., 2022). Due to human error however, the data were instead made unavailable (“kill” command), i.e. made unavailable on the SRA (Berman *et al*., 2022). The avoidable deletion of raw data motivated speculation that caused harm to scientists who had submitted them.

While Bloom (2021b) acknowledged that information was still available in Wang *et al*. (2020b)’s Table 1, he argued that “*the practical consequence of removing the data from the SRA was that nobody was aware these sequences existed*”. However, phylogenetic analyses of SARS-CoV-2 including that performed in Bloom (2021b) routinely exclude sequences with much higher coverage than any sample in this dataset. We are unaware of additional phylogenetic studies in the last 3 years that have utilized Wang *et al*. (2020b)’s data to draw new conclusions about early SARS-CoV-2.

Although of limited value for phylogenetic studies because of their very partial coverage, the data shared by (Wang *et al*., 2020b) are not uninformative. The mutations reported in Wang *et al*. (2020b)’s Table 1 include additional mutations from early SARS-CoV-2 lineages, including one mutation (A24325G, sample B9) found in a sample from a Huanan market vendor (World Health Organization, 2021). Two mutations are also found in two samples each that characterize sublineages derived from lineage B (G22081A, samples C1 and D11) and lineage A (C24034T, samples E5 and R15), indicating that these are not samples from patients with the earliest infections of the pandemic.

In Bloom (2021b), the removal of the raw data was presented as part of a larger narrative, set up in his Introduction, in which Chinese authorities would have gagged researchers and made them retract or falsely amend previous statements, in particular on early Covid-19 cases. In this narrative, changes in the inclusion of early cases are seen as censorship rather than the simple correction of errors. The censorship narrative comes from the fanciful interpretation of a news article published on a blog by a lab leak activist.^19^ The original news article,^20^ however, does not support this narrative, when the quotes are read in full (emphasis added):

> *As of February 25, our entire database has about 47,000 cases. The database has some data on patients who developed the disease before December 8 last year*, ***but we cannotbesureof theauthenticity of these data andfurtherverification is needed***.. *Professor Yu Chuanhua explains*,
>
> *“For example, there is data on a patient who developed the disease on September 29, the data shows that the patient did not undergo nucleic acid testing, the clinical diagnosis (CT diagnosis) is a suspected case and the patient has died, this data has no confirmed diagnosis and no time of death*, ***it could also be wrong data***.*”*

These quotes in the original news article make it clear that the retrospective search of Covid-19 cases was work in progress, and that the results could change. There is no evidence that the researcher was forced by Chinese authorities to walk back earlier comments because of a gag order; instead, the researcher reported on an ongoing analysis of suspected Covid-19 cases. Likewise, it is important to emphasize that the order to destroy samples, mentioned in Bloom’s Introduction, was not specific to Covid-19. Rather, it followed from a biosafety regulation published long before Covid-19.^21^

The notion that Wang *et al*.‘s data withdrawal was linked to something nefarious was pervasive throughout Bloom’s article. In the final version of his article, Bloom (2021b) added a note suggesting that Wang *et al*. might have wanted to retract their preprint to cover their tracks.^22^ There is no evidence that Wang *et al*. wanted to delete their paper. On the contrary, their work was featured in official channels, both before and after the removal of data on SRA (see Table 1). First, the preprint did not contain a link to the data—the data were submitted to the SRA only after the preprint was posted. Second, a press release about the work was posted before the preprint was submitted to medRxiv and was still online at the end of 2021. Third, the peerreviewed paper itself was also advertised: Wuhan University tweeted about the paper when it came out. Finally, the work appears to have been part of a patent application.

Beyond the data from Wang *et al*. (2020b), but still in the context of the Bloom (2021a,b) study, Bloom also investigated other sequence datasets that were either removed or corrected to answer questions about whether sequencing (meta)data relevant to the origin of SARS-CoV-2 had been suppressed. We show in the Appendix that in all cases investigated, the answer is “no.” While we reject unfounded speculation of this sort, we recognize that some datasets relevant to the origin and early spread of SARS-CoV-2 are known to exist and remain unpublished (Holmes, 2024). We hope that these datasets will be published to help resolve unanswered questions.

Bloom’s article illustrates how prejudices can influence scientific conclusions. Data and analysis were presented through the lens of Chinese censorship and the implication that research in China is inherently untrustworthy. We conclude by noting that we initially took it for granted when we read Bloom (2021a)’s claim, in his preprint, that “*the sequences were deleted to obscure their existence*” (ZH), or were initially captivated by the feat of recovering deleted data (FD). The fact that this narrative captured so much attention despite *a complete lack of supporting evidence* prompts us to reflect on how our biases shape our interpretation of data, and how extreme differences in believing people based on where they work can lead to incorrect and harmful conclusions. Here, we are reflecting on our experiences, and we invite readers to do the same.

## Data availability

Data and code are available on Zenodo https://zenodo.org/doi/10.5281/zenodo.10665464.

## Methods

We followed the same methods as Bloom (2021b) to compare sequences to outgroups. We used data shared by Bloom on Github. For the expanded dataset, we used outputs of a dataset curated by Pekar *et al*. (2022), complemented by data recently shared by Lv *et al*. (2024) (selecting the earliest informative sequence for each patient; Genbank accessions are listed in the associated Zenodo repository). We gratefully acknowledge the authors from the originating laboratories and the submitting laboratories, who generated and shared through GISAID the viral genomic sequences and metadata on which this research is based. GISAID accessions used are the same as Pekar *et al*. (2022) data S1. The Yan *et al*. (2021) data correspond to EPI_ISL_493149 to EPI_ISL_493190.

## Acknowledgements

FD thanks the SRA team for their answers to her questions. We thank Alex Crits-Christoph, Joel Wertheim and Mike Worobey for comments and discussions. Zhihua Chen first spotted the two Guangdong clusters error in Bloom (2021b). FD thanks Dake Kang for providing details on the Chinese law behind the destruction of samples. FD thanks Wiley for sharing details on the Wang *et al*. (2020b) publication process and on their investigation. We thank Jonathan Pekar and Alex Crits-Christoph for sharing their files of substitutions. Finally, we thank all data producers for sharing their sequencing data on GISAID and on open platforms (the GISAID accessions are those from Pekar *et al*. (2022), listed in their data S1). ZH is supported by Fundação para a Ciência e a Tecnologia (FCT) through MOSTMICRO-ITQB (UIDB/04612/2020, UIDP/04612/2020) and LS4FUTURE (LA/P/0087/2020).

## Appendix

Here we provide details or other datasets related to Bloom’s study. These datasets were cited in different versions of the study or in accompanying communication. They correspond to data that were removed at some point from public repositories, or associated with metadata that changed. In all three cases that we present, the (meta)data do not shed light on the origin of SARS-CoV-2, and the explanations for their removal or modification are mundane.

### SRR11119760 and SRR11119761, PRJNA607174

Bloom’s original preprint (Bloom, 2021a, v1; Figure 2) contained the screenshot of an email showing another group of Chinese scientists asking SRA for the removal of their data. The screenshot had been obtained through a FOIA request by an activist group pursuing the hypothesis that papers on pangolin viruses were part of a concerted diversion.^23^ By pure happenstance, the data were put back online on June 16, 2021,^24^ that is, two days before Bloom posted his preprint to bioRxiv and shared it with NIH leadership, and six days before the preprint was published on bioRxiv (June 22, 2021). The data were then synchronized on the cloud on June 18, 2021, the day Bloom submitted his preprint to bioRxiv and shared it with NIH officials. Consideration of the exact times of the different events on June 18 further confirms that the events are not related. The preprint PDF was indeed generated at 17:52 EST, ^25^ which corresponds to the time of the last Github commit on the preprint’s associated repository.^26^ Communication of the results to NCBI/NIH officials took place at 19:00 EST.^27^ SRR11119760 was however public on the cloud at 14:00 EST,^28^ i.e. before the preprint’s final version was compiled.

The SRA team indicated that the data had be released “*at the request of a user*”. Whether this is related or not, we can note that a Zenodo document was updated on June 21, 2021 with an analysis of that dataset (Daoyu Zhang, 2020, version 14), i.e., before Bloom’s preprint was even published on bioRxiv and therefore before attention was drawn to those data.

#### PRJNA637497

In the revised version of his study, Bloom included an email by SRA confirming that two Wang *et al*. datasets had been removed (Bloom, 2021b, Figure 6). Although the accessions were redacted in Bloom’s article, Bloom shared the second accession on social media,^29^ and the email is available in documents posted on the Internet. The metadata of this dataset are back online on SRA (under SAMN15143806^30^/SRR11931188^31^), and indicate that it contained a single sample collected on 23 March 2020, i.e. too late to be relevant to the origin of SARS-CoV-2. The SRA team confirmed that the collection date has not been modified since the initial submission in 2020.

#### PRJNA605907

A separate study on early cases (Shen *et al*., 2020) was discussed by Bloom in Twitter threads^32^ related to his 2021 MBE article. The main text of the Shen *et al*. article initially was not consistent with sequence metadata; a correction was published after Bloom’s initial tweets (Shen *et al*., 2021). An in-depth examination of the data indicates that the samples were collected as announced in the sequence metadata and as later corrected. The samples were collected from known patients from 30 December 2019, and sent to separate groups for analysis. The patients are known, the timeline is clear, and there is zero evidence that the samples were collected earlier than the stated date.^33^

## Supplementary figures and tables

**Figure S1:**
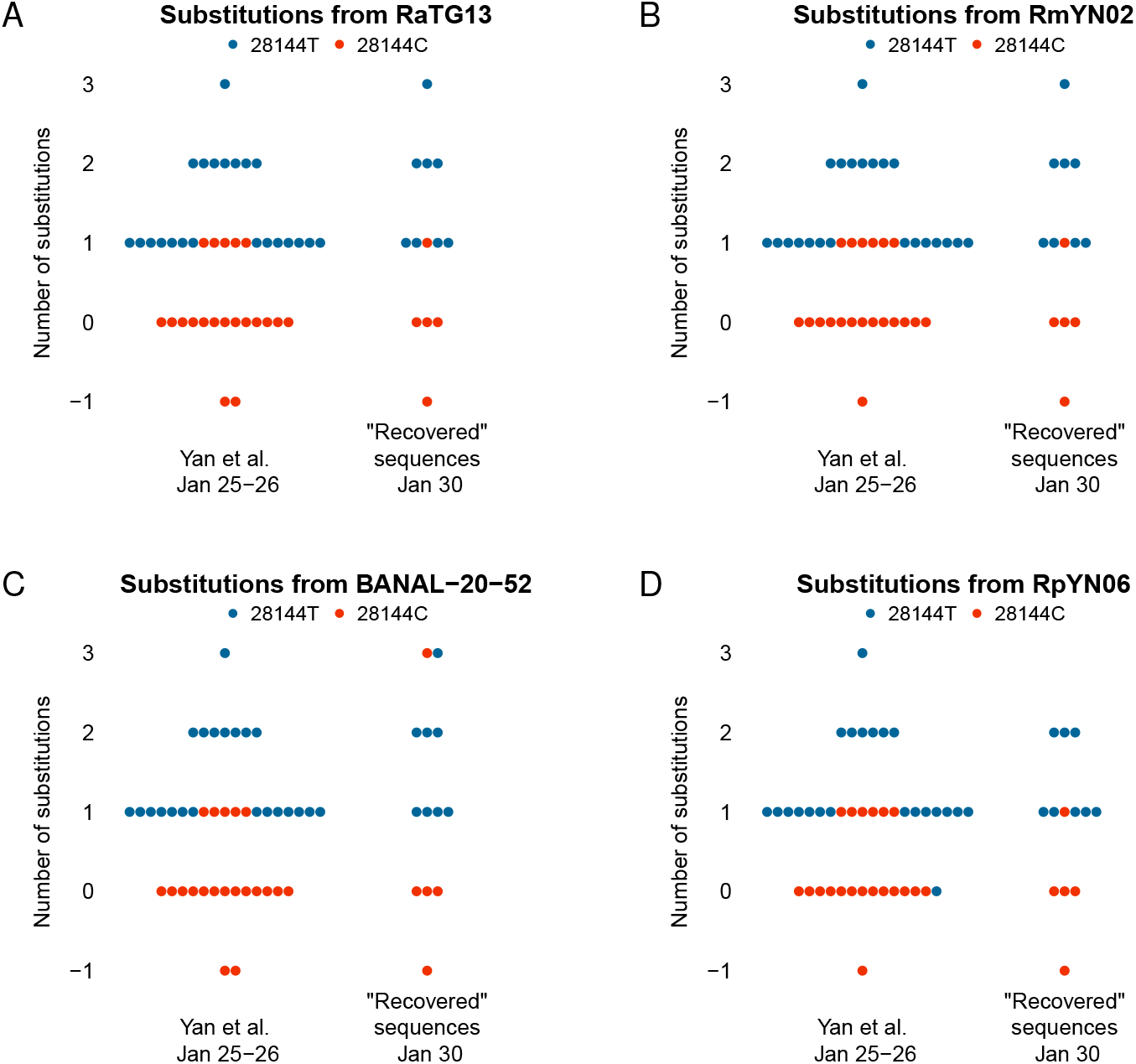
Equivalent of Figure 2, changing the outgroup comparator, shown as title of each panel.

**Figure S2:**
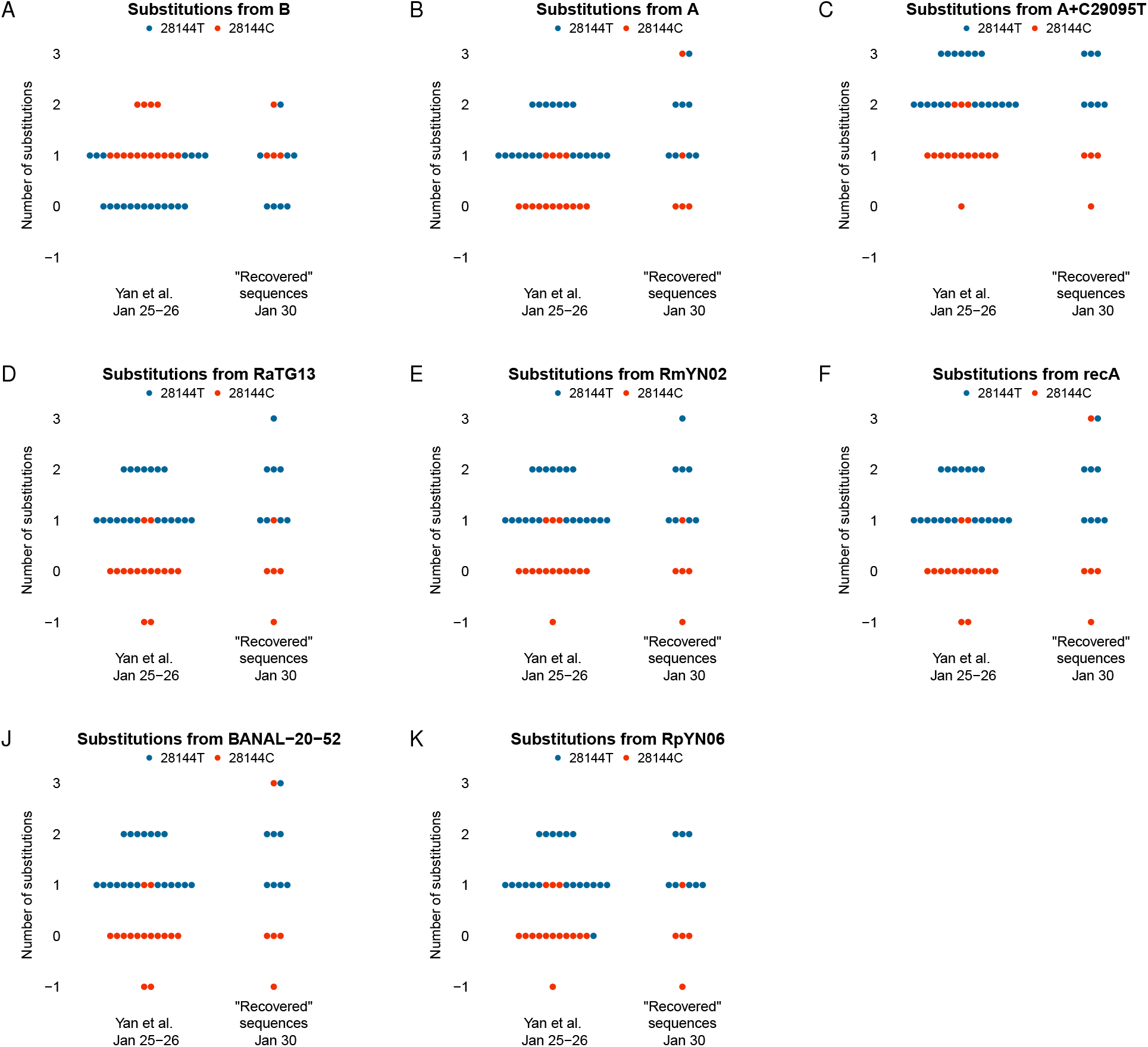
Equivalent of Figure 2, removing sequences with potential sequencing errors (Pekar *et al*. (2022) dataset), and changing the outgroup comparator, shown as title of each panel.

**Figure S3:**
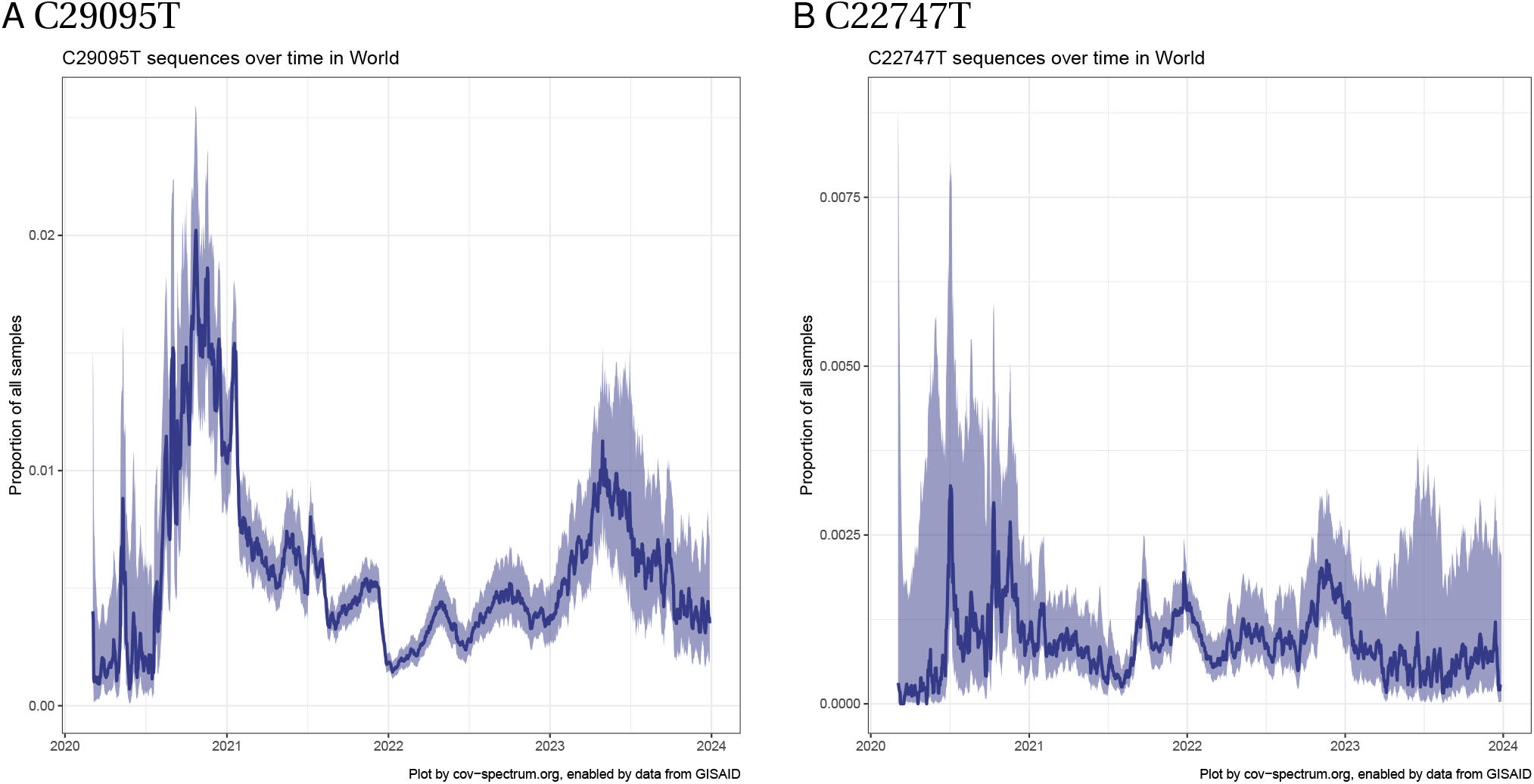
Proportion of sequences with C29095T and with C22747T among all sequences available on GISAID from 1 March 2020 through the end of 2023. Plots generated by CoV-Spectrum (Chen *et al*., 2022), from https://cov-spectrum.org/explore/World/AllSamples/from%3D2020-03-01%26to%3D2024-01-01/variants?nucMutations=C29095T& and https://cov-spectrum.org/explore/World/AllSamples/from%3D2020-03-01%26to%3D2024-01-01/variants?nucMutations=C22747T&. Note the different vertical axis scales.

**Figure S4:**
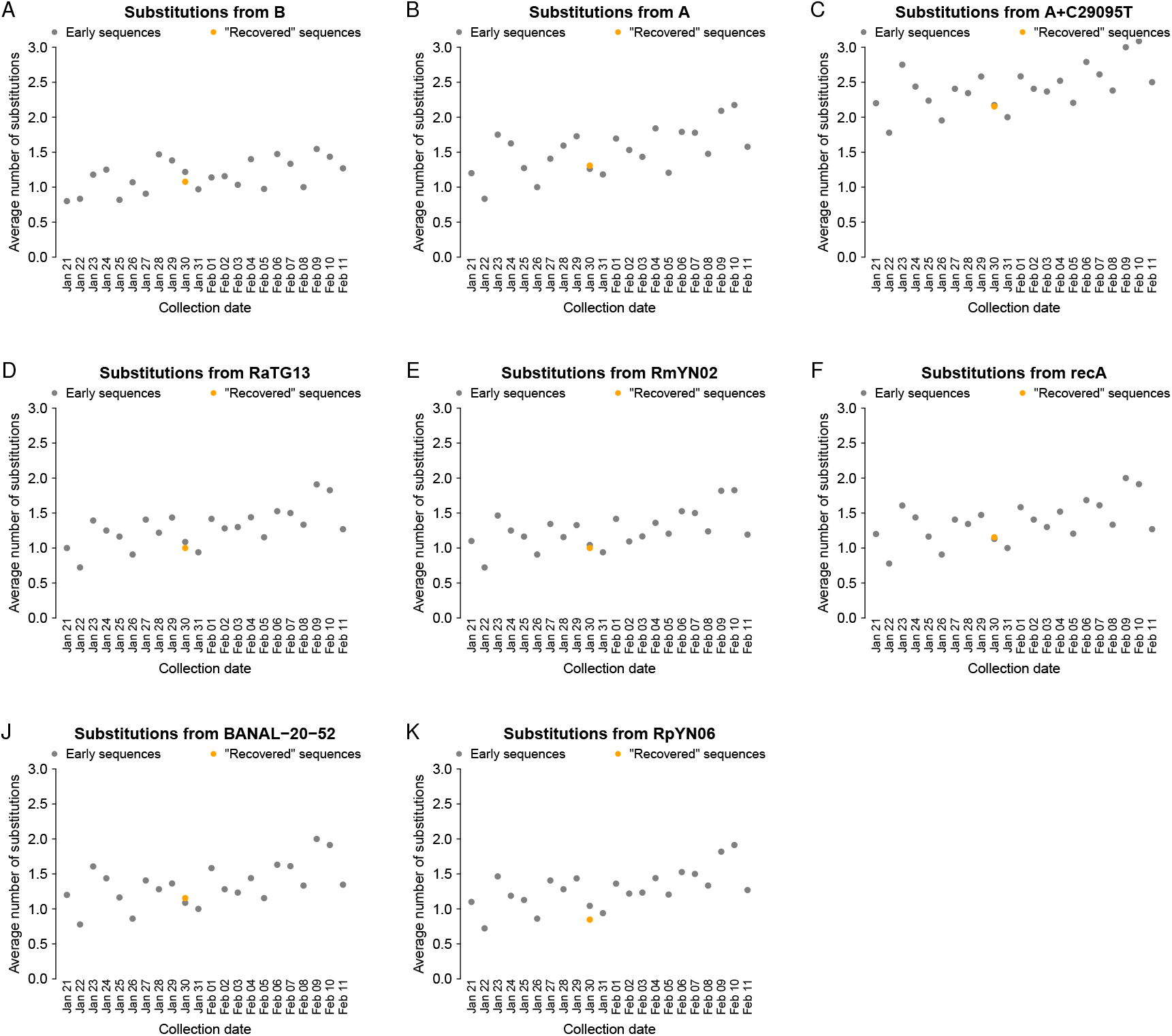
Equivalent of Figure 3, changing the outgroup comparator (shown as panel title).

**Figure S5:**
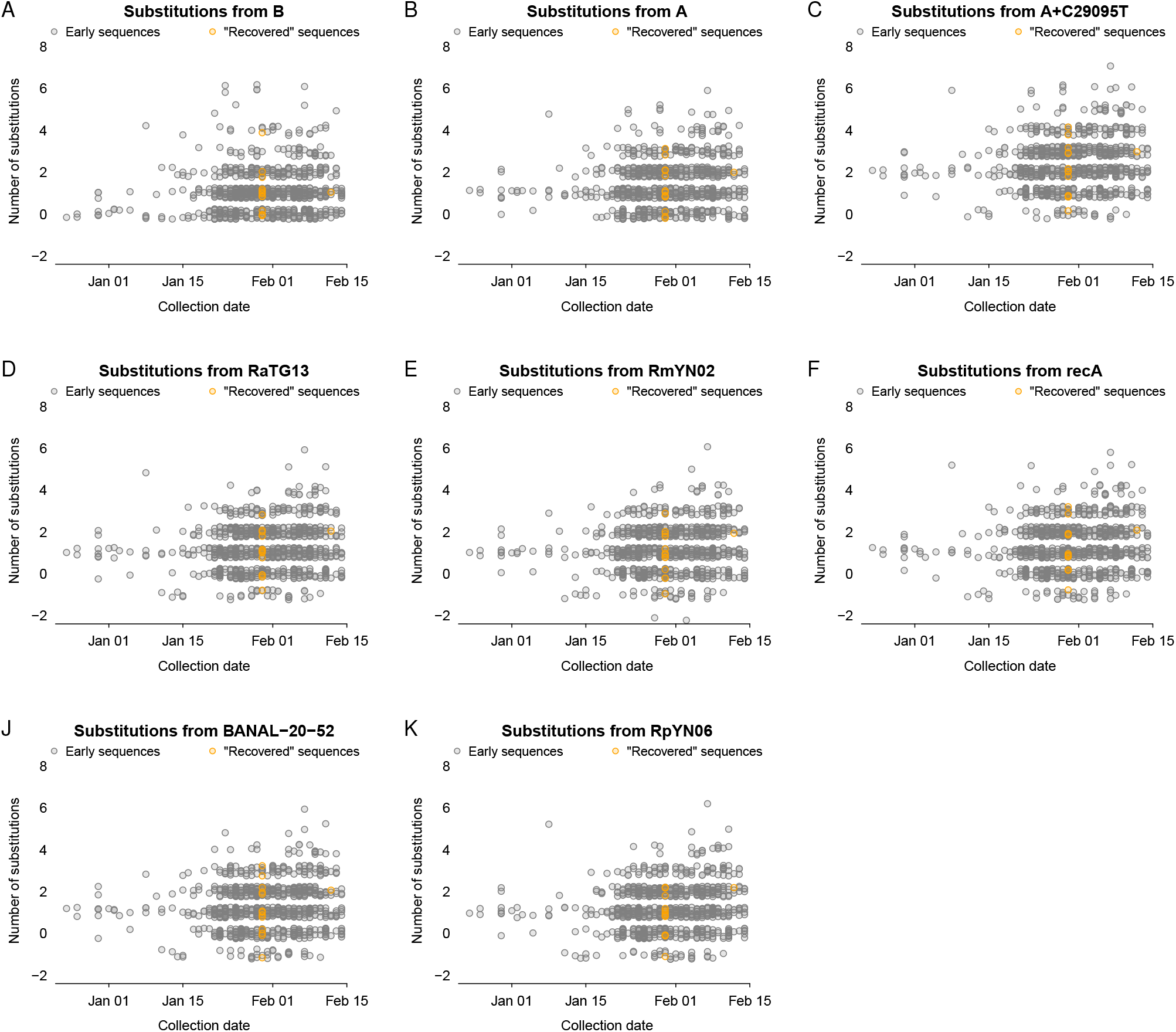
Equivalent of Figure 3, showing all points instead of averages and over a wider time window. The outgroup comparators are shown as panel titles.

**Figure S6:**
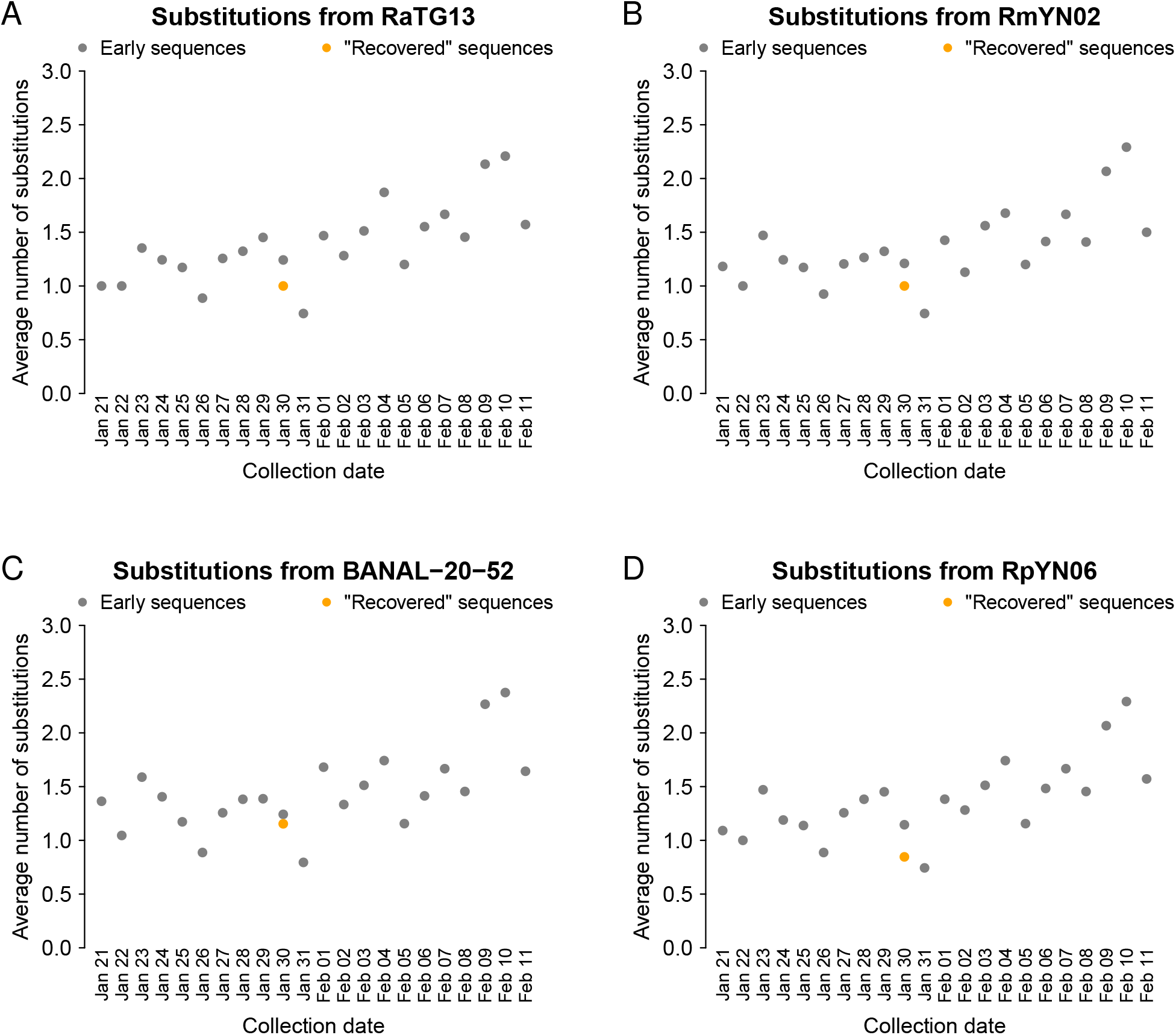
Equivalent of Figure S4, with Bloom (2021b)’s dataset instead of the expanded curated dataset.

**Table S1:** Patients in the early January 2020 Shenzhen cluster. Hospitalization refers to the date of admission at HKU-SZH. Metadata from Chan *et al*. (2020).

| ID | Age | Sex | Link | Onset | Hospitalization |
| --- | --- | --- | --- | --- | --- |
| SZ01 | 65 | F | Mother of SH03 | 2020-01-03 | 2020-01-10 |
| SZ02 | 66 | M | Father of SH03 | 2020-01-04 | 2020-01-10 |
| SZ03 | 37 | F | Daughter of SH01 and SH02 | 2020-01-09 | 2020-01-11 |
| SZ04 | 36 | M | Son in law of SH01 and SH02 | 2020-01-05 | 2020-01-11 |
| SZ05 | 10 | M | Grandson of SH01 and SH02 |  | 2020-01-11 |
| SZ06 | 63 | F | Mother of SH04 |  | 2020-01-11 |

**Table S2:** Sequences in the early January 2020 Shenzhen cluster. Yang *et al*. are Yang Yang, Chenguang Shen, Li Xing, Zhixiang Xu, Haixia Zheng, Yingxia Liu, as listed on GISAID; we have not found a specific article presenting the sequences. The ID column corresponds to patient IDs introduced in Table S1.

| Accession GISAID | ID | Collection | Source | Bloom label |
| --- | --- | --- | --- | --- |
| EPI_ISL_406592 | SZ01 | 2020-01-13 | Yang *et al.* |  |
| EPI_ISL_403933 | SZ01 | 2020-01-15 | Kang *et al.* (2020) | Guangdong patients |
| EPI_ISL_406030 | SZ02 | 2020-01-10 | Chan *et al.* (2020) | Guangdong patients |
| EPI_ISL_406593 | SZ02 | 2020-01-13 | Yang *et al.* | other China |
| EPI_ISL_403932 | SZ02 | 2020-01-14 | Kang *et al.* (2020) | Guangdong patients |
| EPI_ISL_405839 | SZ05 | 2020-01-11 | Chan *et al.* (2020) | other China |
| EPI_ISL_403935 | SZ06 | 2020-01-15 | Kang *et al.* (2020) | Guangdong patients |

## Footnotes

1 References to non-academic work are presented as footnotes.

2 e.g., C. Zimmer, Scientist Finds Early Virus Sequences That Had Been Mysteriously Deleted, https://www.nytimes.com/2021/06/23/science/coronavirus-sequences.html; see https://medrxiv.altmetric.com/details/108029569/news/page:3 for other examples.

3 Press conference recording, 22 July 2021: https://www.youtube.com/watch?v=UA2P8hlurlQ&t=4606s; Transcript: https://www.pekingnology.com/p/why-did-wuhan-university-researchers.

4 There were multiple such official notices at different dates in early 2020; see timeline in Table 1. Different censorship narratives inconsistently refer to one or the other.

5 This dataset is described at https://blog-assets-lifebit.s3-eu-west-1.amazonaws.com/SARS-CoV-2+Public+Dataset.pdf and sample metadata can be found at https://lifebit-sars-cov-2.s3-eu-west-1.amazonaws.com/reads/SraRunTable.txt. Bloom (2021b)’s Materials and Methods section includes a link to an index of URLs for sample data and metadata.

6 Press conference – Zeng Yixin, vice minister of China’s National Health Commission: “*According to our understanding, the earliest sampling time of this batch of samples was January 30 some time has passed since the COVID outbreak began. In fact, it is not an early sample. These sequences provide limited information and value for COVID-19 origin tracing*.”; Zichen Wang, 22 July 2021, Why did Wuhan University researchers delete Covid-19 data at NIH?: https://www.pekingnology.com/p/why-did-wuhan-university-researchers.

7 ”*According to the researcher, a total of two batches of samples were taken. In the first batch, a total of 45 samples were taken randomly from patients that sought treatment in Wuhan on Jan. 30th, 2020. The second batch of samples was taken from a group of patients in mid-February, 2020*.” Zichen Wang, 24 July 2021, The Chinese side of the COVID data withdrawal controversy: https://www.pekingnology.com/p/the-chinese-side-of-the-covid-data (Interview conducted by Yang Liu).

8 Zichen Wang, 24 July 2021, *ibid*.

9 ”*Whenwesawthatthejournalhaddeletedtheparagraph, webelievedthatthentheparagraphwasunnecessary*.”, *ibid*.

10 ”*Because the paper no longer included this descriptive paragraph (of the link to the database), the data that was stored in the database was like a headless fly. Nobody would know the data’s association, maybe after some time, even we wouldn’t be able to find the data, since there was no link. So we asked for the data to be deleted. This took place in June 2020*.”, *ibid*.

11 Email to FD, 9 February 2024.

12 e.g., https://www.ncbi.nlm.nih.gov/biosample/?term=SAMN14381071.

13 hCoV-19/Wuhan/0126-C13/2020 in the A + C18060T root, and hCoV-19/Wuhan/0126-C31/2020 in the A + C29095T root. The C31 sequence has two additional mutations, but they are unique mutations in Bloom (2021b)’s dataset and were therefore discarded in his phylogenetic analysis.

14 We note that “half its progenitor node” was true in Bloom (2021a), but is not in Bloom (2021b), owing to a shift in methods from suppressing rare haplotypes to suppressing rare mutations.

15 Considering full sequences in Bloom’s data set collected on Jan 30, 2020 *±*5 days, those with C29095T have, on average, 1.1 substitutions (*±*0.2; *n* = 27) outside of the region covered by the “recovered” sequences.

16 Hubei Daily, 27 January 2020: https://news.cri.cn/20200127/615de051-1480-73bf-1d61-7385deaa3ca6.html

17 e.g. [emphasis added], “*Details of the genetic makeup of* ***some of the earliest samples*** *of coronavirus in China were removed from an American database where they were initially stored at the request of Chinese researchers, U.S. officials confirmed, adding to concerns over secrecy surrounding the outbreak and its origins*.”, https://www.bloomberg.com/news/articles/2021-06-24/u-s-confirms-removal-of-wuhan-virus-sequences-from-database; “*Chinese scientists have deleted crucial data* ***from the earliest confirmed Covid patients***, *it emerged today amid intense scrutiny about the true origins of the disease*.” https://www.dailymail.co.uk/news/article-9716531/More-proof-support-lab-leak-theory-China-DELETED-samples-earliest-patients.html; “*Details of the genetic makeup of* ***some of the earliest samples*** *of coronavirus in China were removed from an American database where they were initially stored at the request of Chinese researchers, U.S. officials confirmed, adding to concerns over secrecy surrounding the outbreak and its origins*.”, https://fortune.com/2021/06/24/coronavirus-sample-us-database-removal-china-request/.

18 Zichen Wang, 22 July 2021, *ibid*.

19 While this provenance is not explicitly acknowledged, a Google Drive page in Bloom (2021b)’s reference list points to the blog post, which is also listed in reading notes on Bloom’s Github repository. The blog post contains the same error as in Bloom’s text (namely, mixing up Chinese family names and first names, i.e., writing “Professor Chuanhua”, instead of “Professor Yu”).

20 Health Times, 2020, quote from translation referenced in Bloom (2021b) of article available online in syndication at https://www.guancha.cn/politics/2020_02_27_538822.shtml; a truncated article published in print is available at http://paper.people.com.cn/jksb/html/2020-03/03/content_1974813.htm.

21 Law text: https://www.gov.cn/gongbao/content/2019/content_5468882.htm.

22 ”*Notably, it is not possible to delete preprints from bioRxiv and medRxiv, so once Wang* et al. *(2020) had posted their preprint, it was permanently committed to the public record (withdrawn preprints are still accessible, for instance see Yang et al. 2020)*.”, Bloom (2021b).

23 https://usrtk.org/wp-content/uploads/2020/12/NCBI-Emails.pdf.

24 See “Published date”, https://www.ncbi.nlm.nih.gov/sra/?term=SRR11119760 and https://www.ncbi.nlm.nih.gov/sra/?term=SRR11119761.

25 cf. Bloom (2021a)’s pdf metadata.

26 cf. https://github.com/jbloom/SARS-CoV-2_PRJNA612766/commit/cdc5341980c0bf148600ff73bfa1272e7cf09851.

27 according to the time shown on FOIA’s emails, cf. last pages of https://justthenews.com/sites/default/files/2022-03/nih-foia-request-56712_redacted.pdf.

28 as shown with the use of the vdb-dump –info command; https://github.com/jbloom/SARS-CoV-2_PRJNA612766/blob/main/paper/figures/SRR1119760_SRR1119761_obj_timestamps.png.

29 Jesse Bloom, March 31, 2022: “*Finally, e-mails show Wuhan University deleted *two* projects, only one of which (SUB7147304=PRJNA612766) was published in journal Small & described in my paper. Initial emailfocused on deleting another previously unknown project (SUB7554642=PRJNA637497). (23/n)*” https://x.com/jbloom_lab/status/1509598938772361218

30 https://www.ncbi.nlm.nih.gov/biosample/?term=SAMN15143806.

31 https://trace.ncbi.nlm.nih.gov/Traces/?view=run_browser&acc=SRR11931188&display=data-access.

32 https://x.com/jbloom_lab/status/1432903935312818178 on September 1st, 2021 and https://x.com/jbloom_lab/status/1509599601753395210 on March 31, 2022.

33 see https://github.com/flodebarre/Shen-etal_2020/tree/main for an analysis.

